# Trait components of whole plant water use efficiency are defined by unique, environmentally responsive genetic signatures in the model C_4_ grass *Setaria*

**DOI:** 10.1101/234708

**Authors:** Max J. Feldman, Patrick Z. Ellsworth, Noah Fahlgren, Malia A. Gehan, Asaph B. Cousins, Ivan Baxter

**Author notes:** Corresponding author; (IB). One sentence summary: Regulation of plant growth can be partitioned into water dependent and water independent processes controlled by unique genetic components.

## Abstract

Plant growth and water use are interrelated processes influenced by the genetic control of both plant morphological and biochemical characteristics. Improving plant water use efficiency (WUE) to sustain growth in different environments is an important breeding objective that can improve crop yields and enhance agricultural sustainability. However, genetic improvements of WUE using traditional methods have proven difficult due to low throughput and environmental heterogeneity encountered in field settings. To overcome these limitations the study presented here utilizes a high-throughput phenotyping platform to quantify plant size and water use of an interspecific *Setaria italica* x *Setaria viridis* recombinant inbred line population at daily intervals in both well-watered and water-limited conditions. Our findings indicate that measurements of plant size and water use in this system are strongly correlated; therefore, a linear modeling approach was used to partition this relationship into predicted values of plant size given water use and deviations from this relationship at the genotype level. The resulting traits describing plant size, water use and WUE were all heritable and responsive to soil water availability, allowing for a genetic dissection of the components of plant WUE under different watering treatments. Linkage mapping identified major loci underlying two different pleiotropic components of WUE. This study indicates that alleles controlling WUE derived from both wild and domesticated accessions of the model C_4_ species *Setaria* can be utilized to predictably modulate trait values given a specified precipitation regime.

## INTRODUCTION

Improving crop productivity while simultaneously reducing agricultural water input is essential to ensure the security of our global food supply and protect our diminishing fresh water resources. Agriculture is by far the greatest industrial consumer of fresh water, largely because productivity losses related to drought stress can decrease crop yields by greater than 50% (Boyer, 1982; Hamdy et al., 2003). Addressing these challenges will require an integrated approach that combines irrigation practices that minimize water loss and deployment of crop plants with superior water use efficiency (Boutraa, 2010; Davies and Bennett, 2015; Evans and Sadler, 2008; Gregory and George, 2011; Morison et al., 2008; Stanhill, 1986).

Plant water use efficiency (WUE) can be broadly defined as the ratio of biomass produced to total water lost by the plant (Bacon, 2009; Blum, 2009; Condon, 2004; Evans and Sadler, 2008; Monteith, 1993; Morison et al., 2008; Tardieu, 2013). This complex trait is determined by many factors including photosynthetic carbon assimilated per unit of water transpired (Condon et al., 2002; Farquhar et al., 1989; Morison et al., 2008; Penman and Schofield, 1951; Seibt et al., 2008), leaf architecture (Brodribb et al., 2007; Sack and Holbrook, 2006), stomata characteristics (Franks and Farquhar, 2006; Lawson and Blatt, 2014), epidermal wax content (Premachandra et al., 1994), canopy and root architecture (White and Snow, 2012; Martre et al., 2001), stomatal dynamics (Blatt, 2000; Hetherington and Woodward, 2003; Lawson et al., 2010; Flood et al., 2011; Lawson et al., 2012), hydraulic transport (Edwards et al., 2012; Holloway-Phillips and Brodribb, 2011), portion of carbon lost from respiration (Escalona et al., 2012; Tomás et al., 2014) and partitioning of photo-assimilate (Carmo-Silva et al., 2009; Chaves, 1991). Given that plant species (Stewart et al., 1995; Winter et al., 2005; Zegada-Lizarazu and Iijima, 2005; Zhou et al., 2012) and ecotypes within species (Kenney et al., 2014; Lopez et al., 2015; Nakhforoosh et al., 2016; Pater et al., 2017; Ryan et al., 2016; Xu et al., 2009) exhibit variation in WUE it is likely that the characteristics which determine this trait are under genetic control and have evolved in response to different environmental conditions such as water availability (Assouline and Or, 2013; Brodribb et al., 2009; Huxman et al., 2004). Therefore, WUE is likely influenced by both genetically encoded developmental programs and changes in growth environments throughout the plant lifecycle (Fleury et al., 2010).

The technical challenges associated with measuring plant size and transpiration in large structured genetic populations has historically limited experimental efforts aimed at identifying the genetic components associated with WUE. This is particularly difficult in field settings due to year-to-year climate fluctuation and micro-environmental variation observed within agricultural fields. The advent of controlled environment, high-throughput phenotyping instruments (Chen et al., 2014; Fahlgren et al., 2015; Granier et al., 2006; Pereyra-Irujo et al., 2012; Reuzeau et al., 2006; Sadok et al., 2007; Tisné et al., 2013; Walter et al., 2007) alleviates many of these challenges through stringent control of climatic variables and automated, high-resolution measurement of plant size and evapotranspiration across large breeding populations.

Evidence from studies conducted on both crop and model plants indicate that the traits associated with WUE are heritable and largely polygenic, although identifying the causal locus associated with differential performance has proven difficult in crop plants due to plant size and genome complexity (Adiredjo et al., 2014; Aparna et al., 2015; Chen et al., 2012; Coupel-Ledru et al., 2016; Honsdorf et al., 2014; Parent et al., 2015; Schoppach et al., 2016; Xu et al., 2009). Utilization of model plants (C_3_ annuals *Arabidopsis thaliana* and *Brachypodium distachyon*) that possess tractable genetic and experimental properties has enabled scientists to identify QTL that contribute to WUE (Des Marais et al., 2016; Easlon et al., 2014; Lowry et al., 2013; Mojica et al., 2016; Vasseur et al., 2014), a few of which have been mapped to causal genes (Ruggiero et al., 2017). Species in the genus *Setaria* also possess many of these desirable qualities and can be used as experimental models to identify genetic components associated with WUE in a C_4_ plant that is closely related evolutionarily to C_4_ crops like maize, sorghum and bioenergy grasses (Bennetzen et al., 2012; Brutnell et al., 2010; Huang et al., 2016; Li and Brutnell, 2011; Zhu et al., 2017). However, in order to study the diversity of resource utilization tactics present in natural and mapping populations of *Setaria* (Saha et al., 2016) or other C_4_ plant species, methods to quantify plant performance and WUE in different environments must be developed.

The objectives of this study were to use a controlled environment high-throughput phenotyping system to characterize the genetic architecture of plant size, water use and WUE in an interspecific *Setaria* recombinant inbred population (RIL) under two different watering regimes. Our findings indicate that plant size, water use and WUE are polygenic traits which are influenced by both soil water content and greater than 10 pleiotropic loci whose effect size changes differentially throughout development. In addition, we identify and discuss several aspects of experimental design that should be considered when performing high-throughput phenomics experiments to study plant WUE.

## MATERIALS AND METHODS

### Plant material and growth conditions

The experiment here was first described in (Feldman et al., 2017), which focused on plant height, and the details are paraphrased here in quotes for clarity. An interspecific *Setaria* F7 RIL population comprised of 189 genotypes (1138 individuals) was used for genetic mapping. The RIL population was generated through a cross between the wild-type green foxtail *S viridis* accession, A10, and the domesticated *S. italica* foxtail millet accession, B100 (Bennetzen et al., 2012; Devos et al., 1998; Wang et al., 1998). After a six-week stratification in moist long fiber sphagnum moss (Luster Leaf Products Inc., USA) at 4°C, *Setaria* seeds were planted in 10 cm diameter white pots pre-filled with ~470 cm3 of Metro-Mix 360 soil (Hummert, USA) and 0.5 g of Osmocote Classic 14-14-14 fertilizer (Everris, USA). After planting, seeds were given seven days to germinate in a Conviron growth chamber with long day photoperiod (16 h day/8 h night; light intensity 230 μmol/m^2^/s) at 31°C day/21°C night before being loaded onto the Bellwether Phenotyping System using a random block design replicating each genotype and treatment combination in triplicate. For each replicate, individual plants of the same genotype were grown side by side with one individual receiving unlimited water supply while the other individual was subjected to water limitation. The growth chamber location of each of these paired replicates was randomly assigned and did not vary during the course of the experiment. Plants were grown on the system for 25 days under long day photoperiod (16 h day/8 h night; light intensity 500 μmol/m^2^/s) with the same temperature regime used during germination. Relative humidity was maintained between 40 - 80 %. Gravimetric estimation of pot weight was performed 2-3 times per day and water was added to maintain soil volumetric water content at either 33% full-capacity (FC) (water-limited) or 100% FC (well-watered) as determined by (Fahlgren et al., 2015). Prescribed soil water content across both treatment blocks was achieved by 15 days after planting (DAP).

The volume of water transpired by individual plants at each pot weighing was calculated as the difference between the measured pot weight and the weight of the pre-filled pot at pot capacity (100% FC) or the difference between current pot weight and the previous weight measurement if no water was added. At the conclusion of each weighing, if the pot weight was below the set point, water was added to the pot to return soil water content back to its target weight. This strategy effectively maintains soil moisture content at a consistent level within both treatment blocks. To evenly establish seedlings before the water limitation treatment began, equal volumes of water (100% FC) were added to all pots for two days after transfer to the system. At 10 DAP, a dry down phase was initiated (no watering) to establish the water-limited treatment block (40% FC) while continuing to maintain a soil water content of 100% FC within the well-watered treatment block.

### Image acquisition and derived measurements

RGB images of individual plants were acquired using a top-view and a side-view cameras at four different angular rotations (0**°**, 90**°** 180**°**, 270**°**) every other day at the Bellwether Phenotyping Facility (Fahlgren et al., 2015). Optical zoom was adjusted throughout the experiment to ensure accurate quantification of traits throughout plant development. The unprocessed images and the details of the configuration settings can be found at the following download site: (https://plantcv.danforthcenter.org/pages/data-sets/setaria_height.html). Plant objects were extracted from images and analyzed using custom PlantCV Python scripts specific to each camera (side-view or top-view), zoom level, and lifter height (https://github.com/maxjfeldman/Feldman_Elsworth_Setaria_WUE_2017). Scaling factors relating pixel area and distance to ground truth measurements calculated by (Fahlgren et al., 2015) were used to translate pixels to relative area (pixels/cm^2^) and relative distance (pixels/cm).

### Biomass estimation

At the conclusion of the experiment, 176 individual plants (91 plants from the 100% FC and 85 from the 40% FC) were harvested to measure aboveground biomass. Gravimetric measurement of fresh weight and saturated fresh weight were taken directly upon tissue harvest after which plant tissue was placed into polypropylene micro-perforated bags (PJP MarketPlace #361001), dried for three days at 60 °C and subsequently weighed to determine dry weight biomass. Multivariate linear regression was used to evaluate, select and calibrate a predictive model to estimate both fresh and dry weight plant biomass.

Regressing plant fresh weight biomass as a function of side-view area, perimeter length, height, object solidity and width indicated that each of these terms is a significant predictor of fresh weight biomass after stepwise model selection using Akaikes Information Criterion (AIC) (Bozdogan, 1987); multiple R^2^ = 0.89). Unlike fresh weight biomass, side-view area, width, and height were the only significant terms used for prediction of dry weight biomass after using the AIC stepwise model selection correction procedure (multiple R^2^ = 0.76). Models containing all significant terms and their interaction achieved a greater model fit, but they introduced artifacts at earlier developmental time points due to model over-fitting (Fig. S1). Generally, models constructed to estimate fresh weight biomass in the well-watered treatment block exhibited greater explanatory power than those constructed to predict dry weight biomass or those in water-limited treatment blocks(Fig.1).

A minimal model containing only the most significant term (side-view area) in both fresh and dry weight models produced a goodness of fit similar to more complex models (fresh weight R^2^ = 0.86; dry weight R^2^ = 0.74). To avoid propagation of error, values that incorporated plant fresh weight biomass were calculated based on adjusted side-view pixel area and translated to fresh weight biomass after analysis. Cumulative biomass values calculated on a genotype within treatment basis were interpolated using loess smoothing (Chambers and Hastie, 1992). Plant size accumulation on a per day basis was calculated as the difference between the loess fit values on a given day and the estimates from the previous day.

### Water loss tabulation

The LemnaTec instrument at the Bellwether Phenotyping Facility provided measurements of water use based upon the gravimetric weight of each pot before watering, the volume of water applied, and the resulting weight after watering. On days when the volume of water added to a pot was greater than zero, the daily volume of water added was the sum of water volume added over a single calendar day. On days when water was not added (e.g. during the dry down period), the water volume was calculated as the minimum gravimetric weight of the pot on the day in question subtracted from the minimum weight value from the previous day. The cumulative volume of water used on a specific day was the sum of all water used prior to that day. By fifteen days after planting (DAP), the dry down phase for the water-limited treatment group was complete and pots containing plants lost substantially more water than their empty pot counterparts in the well-watered treatment group (Fig. 1). This observation indicates pot water loss cannot be considered a proximity measure of total plant transpiration before day 15 in this experiment (Fig. 1). Examination of the ratio between fresh weight biomass accumulated relative to the amount of water used and mathematical prediction of the amount of water used per day over this period suggests that the amount of water used between day 15 and 17 can be used as an approximation of cumulative water transpired by the plant throughout this experiment up to this point (Fig. S2). This data and the observation that at day 17 the plants are still relatively small (less than 8% of their maximum size on average) support the rationale of starting the analysis on this day (Fig. 1). Volumes of water use (daily and cumulative) on a genotype within treatment basis were estimated using loess smoothing.

### Heritability and trait variance partitioning

We used the same approach as in (Feldman et al., 2017) and the details are paraphrased here in quotes for clarity. During this experiment, plant area was measured every other day, so the number of replicates per treatment to calculate broad sense heritability on any given day was limited. To alleviate this technical shortcoming, trait values for each individual were interpolated across missing days using loess smoothing.

Variance components corresponding to broad sense heritability and total variance explained was estimated using a mixed linear model using the R package lme4 (Bates et al., 2015). Broad sense heritability was calculated using two methods. Within an individual experiment, broad sense heritability on a line-estimate basis was calculated using the following formula:

Equation 1:

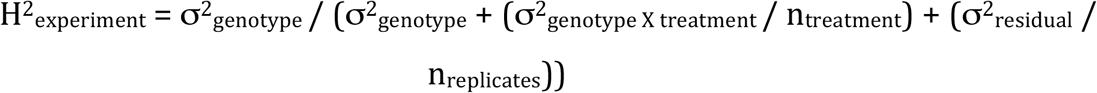

in which n_treatment_ is the harmonic mean of the number of treatment blocks in which each line was observed and n_replicates_ is the harmonic mean of number of replicates of each genotype in the experiment. Heritability within treatment blocks was calculated by fitting a linear model with genotype as the only explanatory factor within each treatment block.

Equation 2:

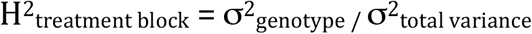

The proportion of variance attributed to genotype divided by total variance within each treatment block is reported as broad sense heritability within treatment (equation). Total variance explained was calculated by fitting a linear model including factors, genotype, treatment, plot and genotype x treatment effects across all phenotypic values in all treatments. The proportion of variance that is incorporated into these factors divided by the total variance in the experiment is reported as total variance explained for each factor.

### QTL analysis

We used the same approach as in (Feldman et al., 2017) and the details are repeated here in quotes for clarity. “QTL mapping was performed at each time point within treatment blocks and on the numerical difference, relative difference and trait ratio calculated between treatment blocks using functions encoded within the R/qtl and funqtl package (Broman et al., 2003; Kwak et al., 2016). The functions were called by a set of custom Python and R scripts (https://github.com/maxjfeldman/foxy_qtl_pipeline). Two complimentary analysis methods were utilized. First, a single QTL model genome scan using Haley-Knott regression was performed to identify QTL exhibiting LOD score peaks greater than a permutation based significance threshold (α = 0.05, n = 1000). Next, a stepwise forward/backward selection procedure was used to identify an additive, multiple QTL model based upon maximization of penalized LOD score. Both procedures were performed at each time point, within treatment blocks and on the numerical difference relative difference and trait ratio calculated between phenotypic values measured in treatment blocks at each time point. QTL associated with difference or ratio composite traits may identify loci associated with genotype by environment interaction (Des Marais et al., 2013).

The function-valued approach described by (Kwak et al., 2016), was used to identify QTL associated with the average (SLOD) and maximum (MLOD) score at each locus throughout the experiment. Each genotypic mean trait within treatments was estimated using a logistic function, and the QTL significance threshold was determined based upon permutation-based likelihood of observing the empirical SLOD or MLOD test statistic. Separate, independent linkage mapping analysis performed at each time point identified a larger number of QTL locations relative to similar function valued analysis based on the SLOD and MLOD statistics calculated at each individual marker throughout the experimental time course. After refinement of QTL position estimates, the significance of fit for the full multiple QTL model was assessed using type III analysis of variance (ANOVA). The contribution of individual loci was assessed using drop-one-term, type III ANOVA. The absolute and relative allelic effect sizes were determined by comparing the fit of the full model to a sub-model with one of the terms removed. All putative protein coding genes (*Setaria viridis* genome version 1.1) found within a 1.5-logarithm of the odds (LOD) confidence interval were reported for each QTL.”

## RESULTS

### Measuring plant size and water use throughout the plant lifecycle

Recurrent measurement of plant size and water use was performed on individuals of a *Setaria* recombinant inbred population (Devos et al., 1998; Devos et al., 1998) grown at two soil water content levels at the Bellwether Phenotyping Facility (Fahlgren et al., 2015). Images of each individual plant were captured every other day from 7 to 33 days after sowing and plant objects were isolated and quantified using PlantCV (Fahlgren et al., 2015; Feldman et al., 2017). Weight estimates of fresh and dry-weight aboveground biomass were calculated using a simple linear model featuring side-view area as the only predictor (Fig. 1, Fig. S3).

**Figure 1.**
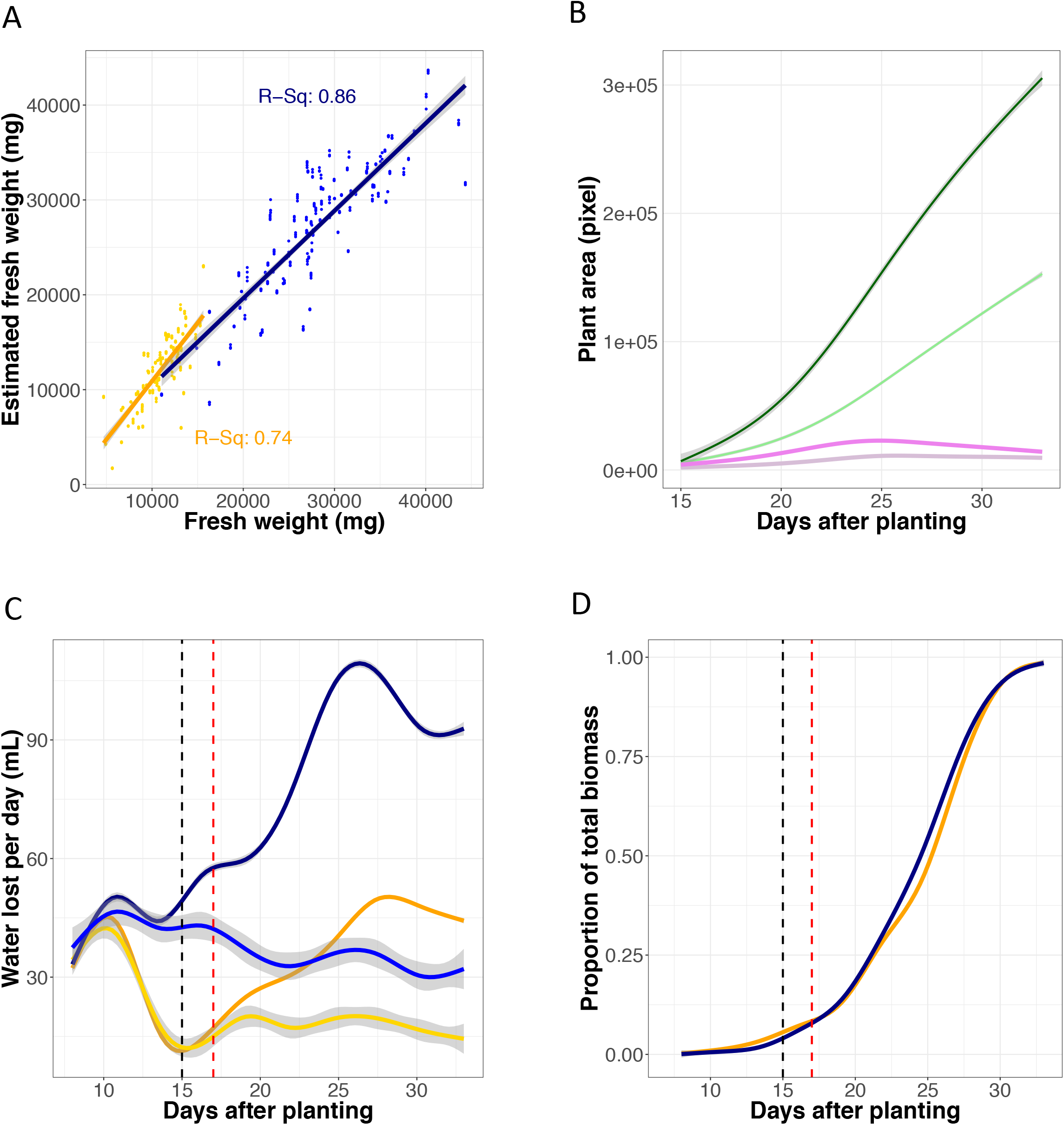
Plant size and water use can be accurately inferred throughout a majority of the plant life cycle. A) Significant correlations between plant fresh weight and pixel area were observed in both the well-watered and water-limited treatment blocks. B) Plants exhibited a sigmoidal growth curve, characterized by an average maximal rate of growth between 23– 26 days after planting. Green lines reflect absolute average size, whereas purple lines report on growth rate. Dark and lighter shaded lines report the well-watered and water-limited treatment blocks respectively. C) Daily water loss can be accurately measured at 17 days after planting. Dark blue and orange lines correspond to average daily water lost from pots, whereas the lines with lighter shades of similar colors report the average water loss of empty pots. The dashed black line denotes the day at which dry down within the water-limited treatment block is complete whereas the dashed red line demarks when water use can be accurately measured. D) By 17 days after planting, plants have attained less than 8% of their total biomass.

Daily plant water use was inferred through gravimetric measurement of pot weight performed two to three times each day by the LemnaTec instrument. The amount of water used by individual plants was calculated as the difference between the measured weight of the pot and the weight of a pre-filled pot at a fixed point that is proportional to its water holding capacity (100% FC) or the difference between current weight and the previous weight measurement if no water was added. At the conclusion of each weighing event, if pot weight was below the set point, water was added to the pot to return it to the target weight value. This strategy effectively maintains soil moisture potential at a consistent level within both treatment blocks. To evenly establish seedlings before the water limitation treatment, equal volumes of water (100% FC) were added to all pots for two days after transfer onto the system. At 10 days after sewing, a dry down phase was initiated (no watering) to establish uniformity within the water-limited treatment block (40% FC) while continuing to maintain a soil water content of 100% FC within the well-watered treatment block.

Examination of water loss from empty pots relative to those containing plants suggested that early in the experiment a majority of water loss was exclusively due to evaporation from the soil surface and did not informatively report on plant transpiration (Fig. 1) (Ge et al., 2016). Beginning the analysis at day 17 enabled us to minimize the artifacts of evaporation that dominated early in the experiment while still capturing growth attributes over a large proportion (~92%) of the plant growth within the experiment (Fig. 1). Another potential confounding issue was the use of a fixed set point for the pot weight, which neglected the increasing weight of the plant when calculating the amount of water needed to return the pot weight to the set point during watering jobs. This decreased the volume of water present within each pot after watering by approximately 12.5% (well-watered) and 17.5% (water-limited) on average by the end of the experiment (Fig.S4).

**Figure 2.**
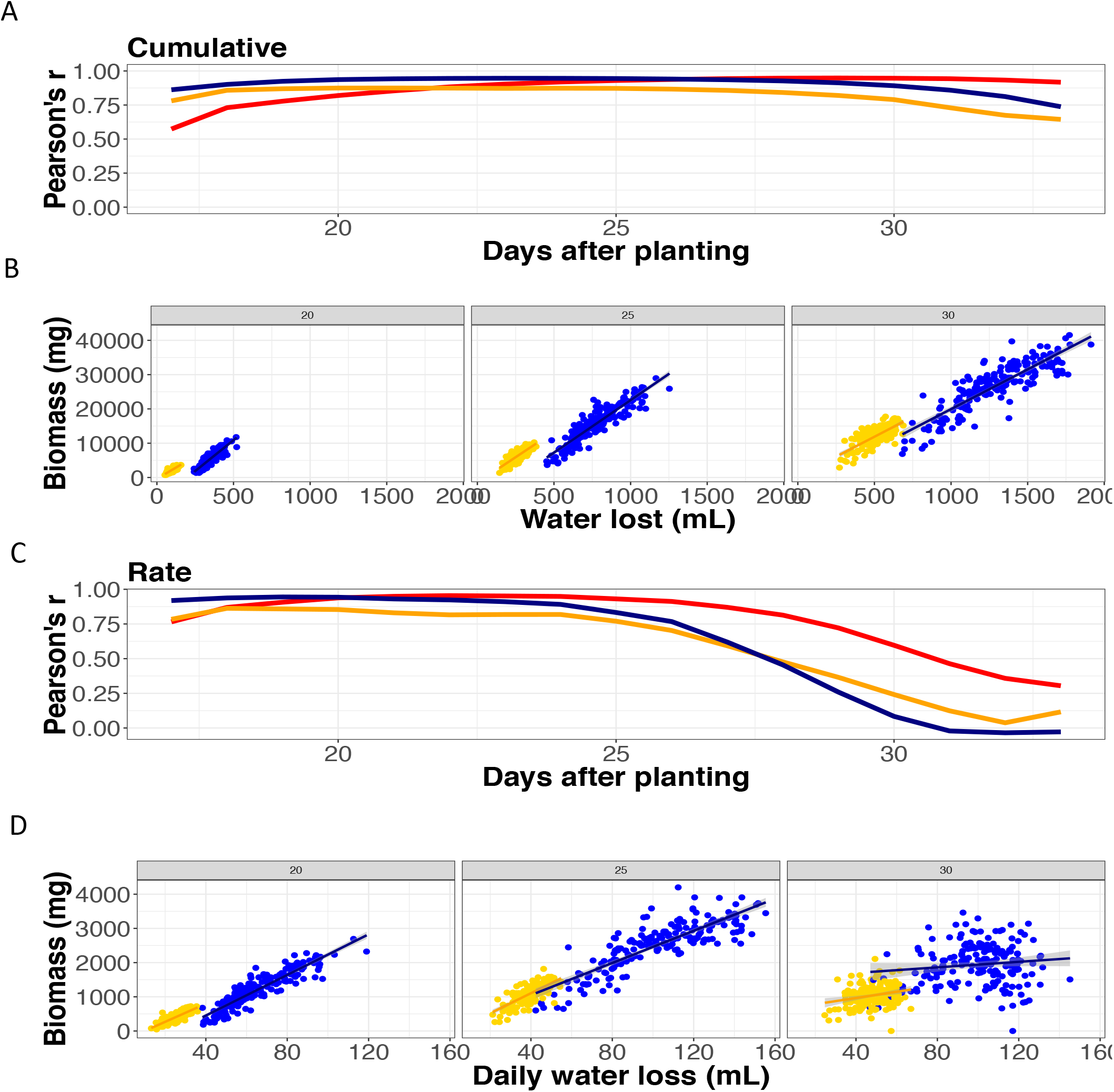
Plant size and water use are tightly correlated. Pearson Correlation Coefficient both within (blue is well-watered, orange is water-limited) and between (red is across both) treatment blocks indicates strong correlation between these two characteristics, although the correlation between the rate of plant growth and daily water use decreases as plants approach maximum size. A) Correlation between cumulative plant size and water use. B) The relationship between plant size and water use at 20, 25 and 30 days after planting. C) Correlation between the rate of plant growth and daily water use. B) The relationship between plant growth rate and daily water use at 20, 25 and 30 days after planting.

Loess smoothing was used to interpolate the values of traits on a genotype level within each treatment block across all experimental time points (Chambers and Hastie, 1992). Fitting of parametric models was avoided because in many cases the trait values exhibited genotype specific temporal responses that could not be accurately represented by a single model type across the entire population. Rate statistics were calculated from these loess smoothed estimates as the difference of the trait between days. Plots illustrating the mean and variance of each trait can be observed in FIG. S5.

### Plant size and water use are correlated

Over the course of this experiment cumulative plant size and water use were highly correlated. Correlation was tightest between 21 and 27 DAP in the well-watered treatment block (> 0.94) and quite strong between 20 and 27 DAP in the water-limited treatment block (> 0.87, Fig. 2). In both treatment blocks, correlations between these characters were weakest at the beginning and end of the experiment but never dropped below 0.67. The correlation of the rate statistics associated with these traits appeared qualitatively different. Correlation between plant growth rate and the rate of water use was initially strong (> 0.79) but rapidly decreased at about 26 DAP as the rate of growth slowed (ultimately approaching zero) by the end of the experiment (Fig. 2) while transpiration remained high.

We implement two numerical approaches to characterize the genetic architecture of the relationship between these traits. The first method, which is hereafter referred to as the water use efficiency ratio (WUE_ratio_), calculated the ratio of biomass relative to the volume of water lost from the pot. This calculation was performed on a cumulative or daily rate basis.

Equation 3:

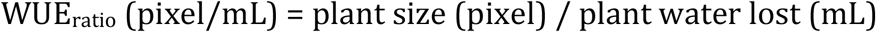

Values of cumulative WUE_ratio_ calculated during this experiment were comparable to other experiments where plant size and water use was measured manually at lower throughput (25-29 grams fresh weight / Liter of water, 7-9 grams dry weight / Liter of water). On average, the cumulative and daily rate WUE_ratio_ was greater in the water-limited treatment block than in well-watered conditions. In principle, the WUE_ratio_ should attenuate the relationship between biomass and water use, but significant correlation was still observed between these two variables, particularly within the rate statistic over the last week of the experiment (Fig. S6, Fig. S7).

**Figure 3.**
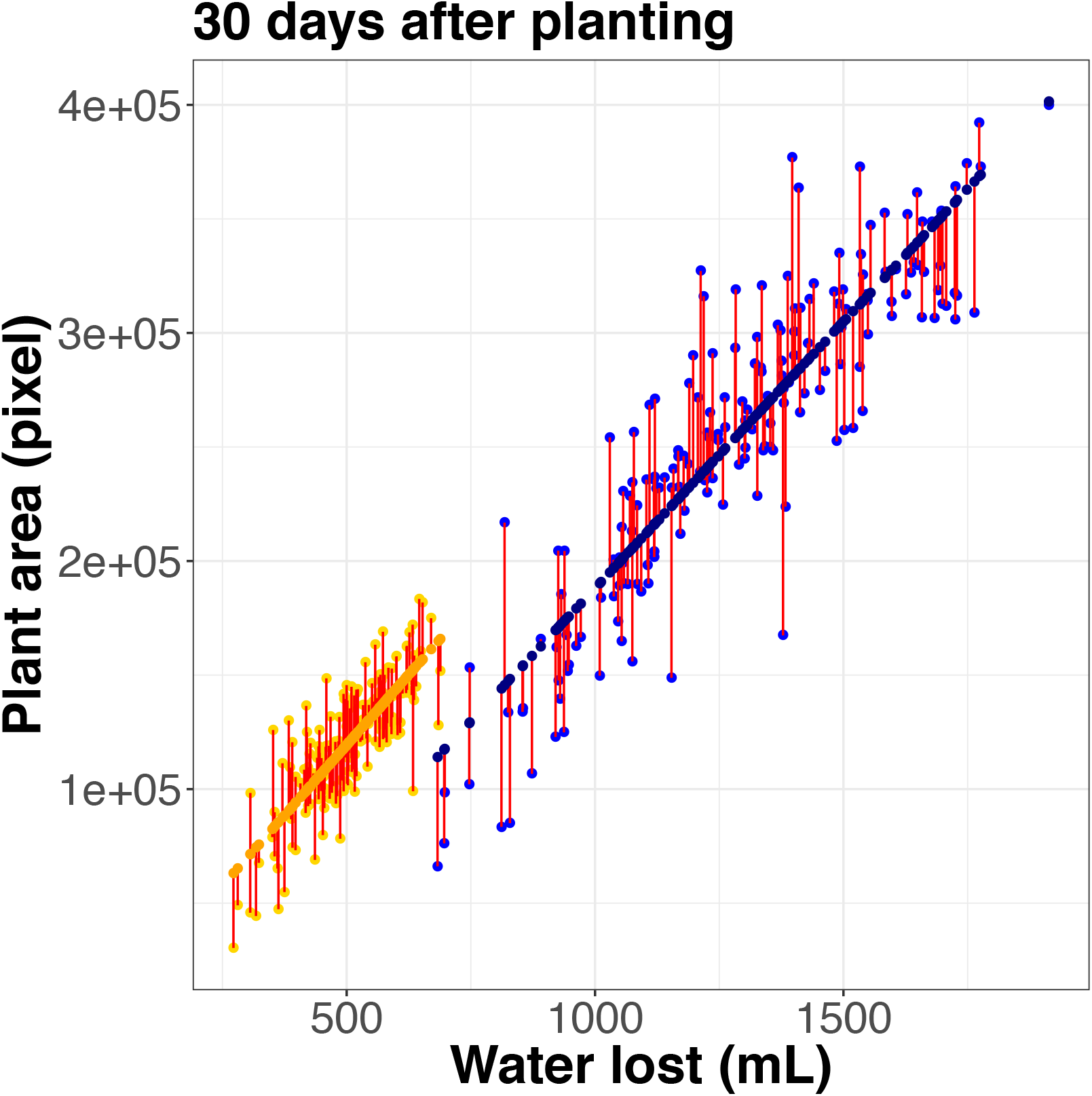
Modeling the relationship between plant size and water use results in two traits. This approach results in predicted value of water use given size (WUE_fit_) colored in dark blue and deviations from this relationship (WUE_residual_) plotted in red. Plot illustrates this relationship at 30 days after planting.

The high correlation between plant size and water use suggests that they were not independent traits in this experimental setup. Therefore, as a second approach, ordinary least squares linear regression was used to model the relationship between plant biomass and water use. For each day of the experiment, within treatment blocks a WUE_model_ was used to predict plant size (dependent/response variable) based upon water loss (independent/explanatory variable) (Fig. 3). The residual of this model fit was evenly distributed around zero across the entire distribution of the predicted values suggesting minimal bias of this approach (Fig. S8).

Equation 4:

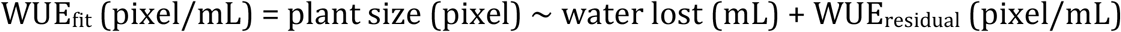

This approach resulted in two traits: The first was the predicted model fit (WUE_fit_) that described the sum of squares relationship between biomass and water use. The residual of this model (WUE_residual_) can be thought of as genotype-specific deviation from this relationship combined with measurement error. As expected, the correlation between the fit values derived from the WUE_model_ was highly correlated with plant size (Fig. S9). A slight correlation between cumulative plant biomass and the residual of the WUE_model_ was observed particularly later in the experiment demonstrating that biomass had components that were not accounted for by the linear model fit (Fig. S10). Varying the dependence structure/assignment or fitting of the model using major axis regression framework (Legendre, 2014) had little effect on downstream analysis.

Each trait (biomass, water loss, WUE_ratio_, WUE_fit_ and WUE_residual_) exhibited high average heritability over all experimental time points within and across treatment blocks (0.28 - 0.77) (Fig. S11). Heritability tended to achieve its maximum value in the middle of the experiment with decreased heritability observed at the beginning and the end of the study. Proportionally, the treatment effect of water limitation explained the largest percentage of variance within biomass, water loss and the WUE_fit_ although genotype and genotype x treatment interaction also explain a substantial margin of the variance (Fig. S12). Heritability of the rate traits was generally similar but on average 5% lower than the heritability of the cumulative traits. In all cases, the average heritability of each trait was greater within the well-watered treatment block relative to the value calculated in water-limited treatment block.

### The genetic architecture of plant size and water use traits

For each day of the experiment, a best fit multiple QTL model was selected for each trait (plant size, water use, WUE_ratio_, WUE_fit_ and WUE_residual_) and the daily rate of change of the trait within each treatment block based upon penalized LOD score using a standard stepwise forward/backward selection procedure (Broman et al., 2003). This approach identified 86 (cumulative Fig. 4; Table S1) and 106 (rate Fig. S13; Table S1) unique SNPs associated with at least one of the five traits. Many of these uniquely identified SNP positions group into clusters of tightly linked loci that are likely representative of a single QTL location. These local clusters of SNPs (10 cM radius) were then condensed into the most significant marker within each cluster to simplify comparisons of genetic architecture between traits (Fig. S13; Fig. S14). Collapsing these SNP positions yielded 23 unique QTL locations associated with cumulative trait values (Fig. 5) and 27 unique rate QTL locations (Table S2).

Of the 23 unique QTL identified, plant biomass contributes the largest proportion of QTL to this set (18) followed by WUE_ratio_ (12), WUE_fit_ (11), WUE_residual_ (10) and water lost (8) (Fig. 5, Fig. S16). Despite the fact that only one QTL location (2@96) was common across all traits and environments, the genetic architecture that contributes to each of these characteristics was clearly related. The strong correlation of plant size and water loss with the predicted value of plant size given water loss (WUE_fit_) are clearly reflected within the genetic architecture associated with these traits. Plant size, water loss and WUE_fit_ all shared 8 QTL (2@96, 3@48, 5@109, 6@65, 7@34, 7@51, 7@99 and 9@34) either within the well-watered or water-limited treatment block (Fig. 5, Fig. S16). Plant size, WUE_ratio_ and deviations from the relationship between plant size and water use (WUE_residual_) shared five QTL unique to this subset (2@11, 2@113, 5@79, 5@92, and 9@127) which enable divergence from the fundamental relationship between plant size and water loss (Fig. 5, Fig. S16). Several QTL were identified as being uniquely associated with plant size (3@21, 5@119, 6@80, 9@138), WUE_residual_ (2@82 3@77, 6@47) and WUE_fit_ (5@39) whereas no QTL were identified as being uniquely associated with water loss or WUE_ratio_ (Fig. 5, Fig. S16).

**Figure 4.**
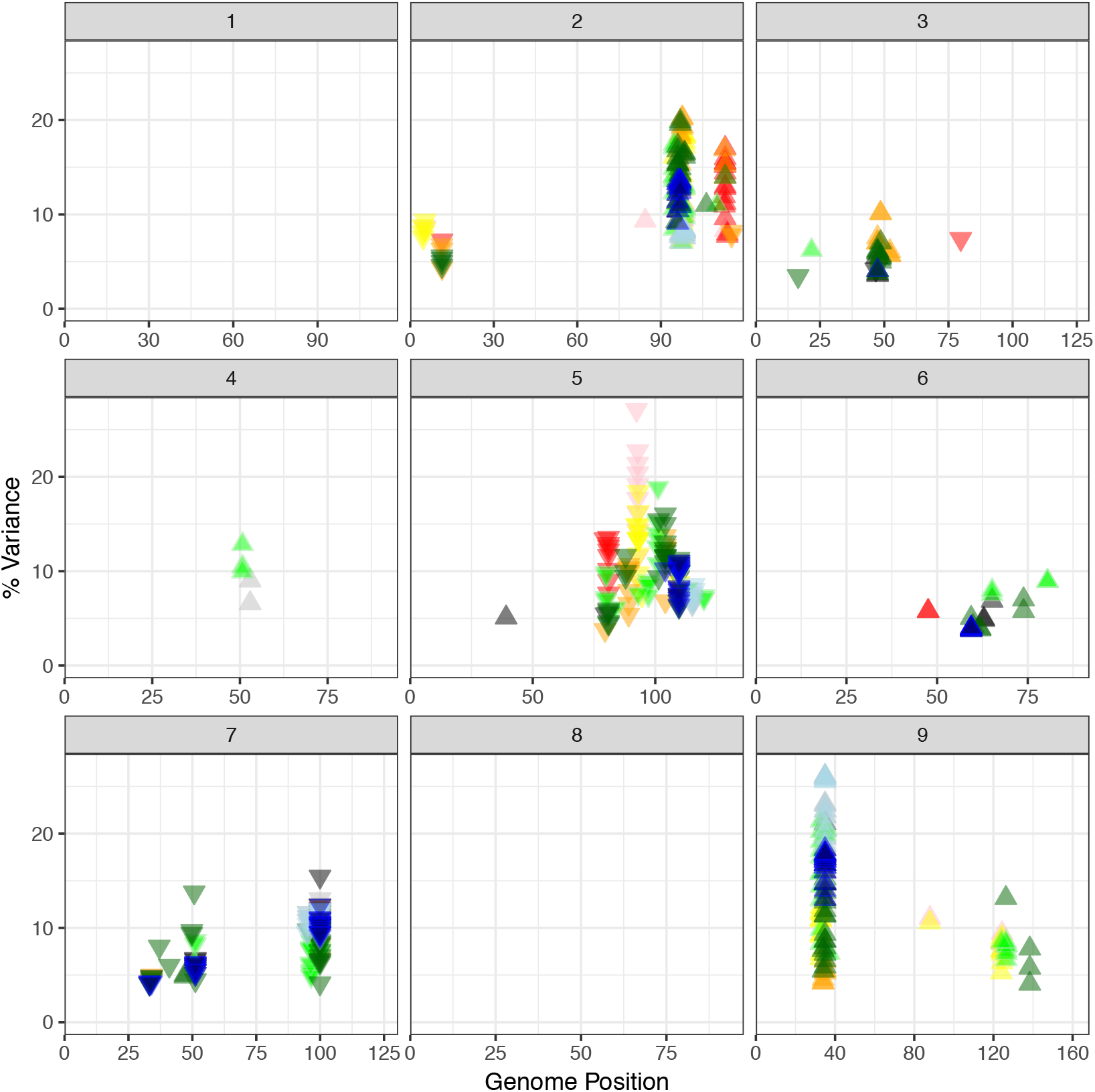
Eighty-six unique QTL locations were detected across all traits in this experiment. Each box corresponds to an individual chromosome, where the values along the x-axis are chromosome position and values along the y-axis denote the proportion of genetic variance explained by the QTL. Each triangle represents a single QTL detected, where the color indicates the trait each QTL is associated with (green = plant size, blue = water use, orange = WUE_ratio_, black = WUE_fit_, red = WUE_residual_). The darkness of color shading is indicative of treatment block where darker represents well-watered and lighter corresponds to the water-limited block respectively. The direction of the arrow indicates the directional effect of the B100 parental allele.

**Figure 5.**
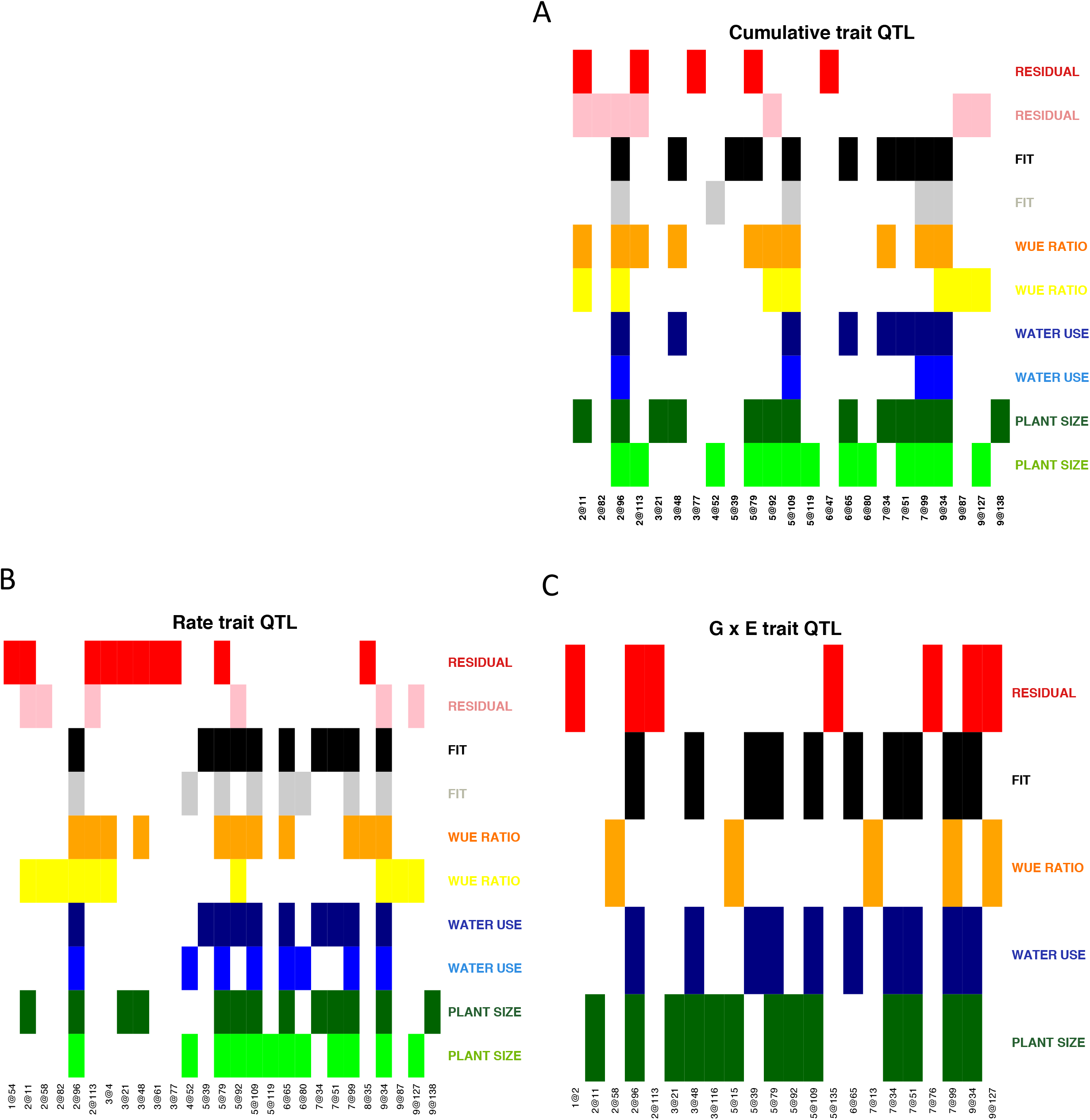
The genetic components that contribute to subsets of traits largely overlap. The QTL locations identified are plotted on the x-axis and the traits are plotted on the y-axis. Colored matrix entries denote at least one significant association within this experiment. A) The genetic architecture of cumulative traits. B) The genetic loci associated with trait rate of change. C) Genetic components associated with genotype x environment traits.

The genetic architecture of all five traits appears to be influenced by water availability. All traits other than water loss exhibited QTL unique to each treatment block (Fig. 5, Fig. S17). Biomass, water lost, and WUE_fit_ all shared four QTL in common across environments (2@96, 5@109, 7@99 and 9@34) where as WUE_ratio_ and the WUE_residual_ shared a single QTL (2@11) between blocks not found associated with the other traits. Two QTL (3@48, 7@34) were found specifically within the well-watered treatment block for all traits other than WUE_residual_ whereas QTL specific to water-limited environment identified common QTL associated with biomass and WUE_fit_ (4@52) or WUE_ratio_ and WUE_residual_ (9@87, 9@127).

The identity of QTL associated with the daily rate values suggest that the genetic architectures were largely cognate with the QTL associated with the traits themselves, both in identity and response to treatment. In total, 28 QTL comprised the union of all unique QTL associated with both the trait value and the daily rate of change calculated from the trait value. Of these QTL, 22 were common between both the trait value and rate statistic associated with the trait, whereas five are only found associated with the rate (1@54, 2@58, 3@4, 3@61, 8@35) and only one QTL was uniquely associated with the cumulative trait values alone (6@47) (Fig. S18).

### Genotype x environment interactions

To assess the genetic architecture of genotype x environment interactions, mapping was performed on numerical difference, relative difference and trait ratio between the phenotypic values observed within each treatment block. In total, 148 unique SNP locations were identified as being significantly associated with at least one of the difference trait formulations across all standard and derived plant size and water use traits (Table S3). Substantial overlap between these categories of genotype x interaction traits indicates that each formulation detects similar genetic signals (Fig. S19) although the large number SNPs found uniquely associated with the trait ratio may indicate that some of these associations maybe spurious. As such, these QTL (trait ratio genotype x environment QTL) were removed from further analysis. The numerical difference and relative difference traits exhibited association with 43 and 40 unique SNP positions, which were representative of 20 and 18 QTL respectively (Table S4, Fig. S20-22).

A majority of the QTL (10/15) identified as being associated with the trait difference between treatment blocks were also found associated with the cumulative trait in both treatment blocks (Fig. 5). The exceptions to this were QTL located on 3@21, 3@48, 5@39, 7@34 and 9@127 that were identified as being significantly associated with the difference between treatment blocks but only identified in either well-watered (3@21, 3@48, 5@39, 7@34) or water-limited conditions (9@127). Interestingly, the QTL located on 3@48, 7@34 and 9@127 were associated with more than one trait in a single treatment block which may indicate that these QTL impart pleiotropic phenotypic effects that were dependent upon soil water content (Fig. 5).

### The temporal genetic architecture of plant growth and water usage

In order to account for the time dependence of the traits for the five plant traits, we used a function-valued approach based upon average log-odds score throughout across the experiment (SLOD) for each trait (Kwak et al., 2016). This analysis parallels the individual time point analysis, although the reduction of complexity (fewer, higher confidence QTL) provides an opportunity for simplification and better understanding of the major loci that influence plant WUE.

**Figure 6.**
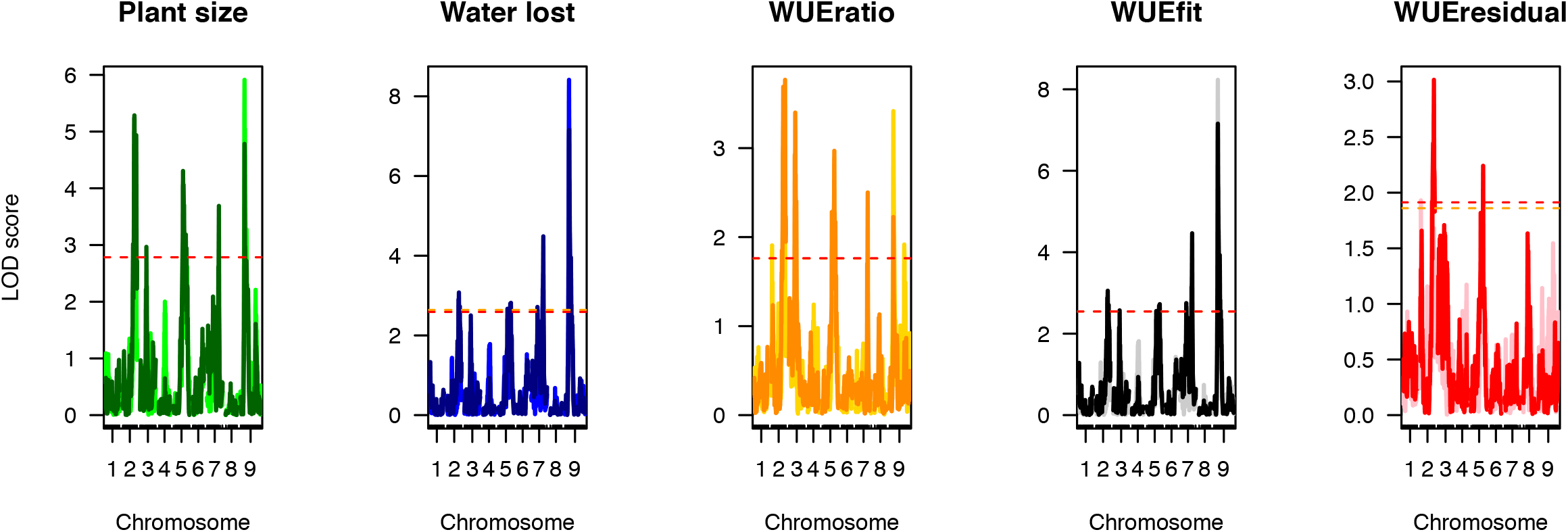
Significant associations identified using single marker scan functional QTL mapping. Chromosomal position is plotted on the x-axis whereas LOD score of trait association across the genome is plotted on the y-axis. Treatment block is indicated by color intensity (darker is well-watered and lighter is water-limited). Significance thresholds (based on 1000 permutations) are plotted as dashed yellow (water-limited) and red (well-watered) lines respectively.

SLOD based function-valued QTL models indicate that several major QTL (2@96, 5@109, 7@99, and9@36) influenced both plant size and water use related traits, although the magnitude of statistical significance attributed to each loci varied by trait and throughout plant development (Fig. 6, Fig S23). Using the SLOD approach, we were able to partition combinations of QTL unique to related traits (Fig. 6). For several QTL (those around 2@96 and 5@109) the positional location at which maximal LOD score was observed changed noticeably in a trait and environment dependent manner either due to multiple closely linked loci or noise in our measurements. Because the confidence intervals of the QTL generally overlap, our reporting in this section will hereafter refer to these loci by their approximate chromosomal location.

Both plant biomass and cumulative water use exhibited almost a complete overlap of QTL within the well-watered treatment block, whereas plant size given water use (WUE_fit_) and deviation of plant size from this fundamental relationship (WUE_residual_) each exhibit a unique genetic signature (Fig 6). As observed when trait values at individual time points were treated as independent traits, a single QTL on 2@96 is the only genetic component that was shared across all five traits. The linear modeling approach successfully partitions out QTL associated with WUE_fit_ (2@96, 7@99, 9@36) from the genetic components that contribute to deviations from the plant size ~ water use relationship (WUE_residual_; 2@96, 5@109). The QTL associated with the WUE_ratio_ (2@96, 3@52, 5@109) also likely reflects deviations from the relationship between biomass given water loss associated with the WUE_residual_. Overall, the identity of QTL associated with each trait was largely identical between the two treatment blocks (Fig. 6, Fig. S23) as were the QTL associated with the values of rate statistics derived from these measurements (Fig. S24, stepwise method; Fig. S25, scanone method).

### A temporal model of the genetic architecture that influences plant water use efficiency

Our QTL results suggest at least two components of water use efficiency with distinct genetic architectures. In order to compare the genetic architecture across all traits, treatments and time points in a common framework, we analyzed how each trait was influenced by a common set of loci. Fourteen QTL were selected based upon their association with multiple traits, robust linkage with a single trait and/or having differential contribution to traits across treatment blocks (Table S5) and the proportional contribution of each locus to the additive genetic variance was calculated using drop-one-term, type III, ANOVA performed for all experimental traits, time points and treatment. Agglomerative hierarchical clustering of the signed proportion of additive genetic variance explained by each locus was performed to identify modules of traits and loci that define plant phenotypes. Examination of scree plots of the within group sum of squares suggested that the variance within traits could be attributed to approximately six groupings although a majority of this variance could be captured within the largest 2-3 partitions (Fig. S26). These partitions represented the major relationships between trait classes. The WUE_ratio_ and WUE_residual_ were generally grouped separately from a larger cluster of traits that included cumulative plant size, water use and WUE_fit_ (Fig. 7). The genetic architecture of plant water use and WUE_fit_ were more related to each other than they were to plant size, which formed the third group. The influence of water availability on these traits was apparent from the grouping of clusters whereas the effects of time were clear but distributed within the treatment blocks. The genetic architecture of the WUE_ratio_ in the well-watered treatment block at early time points was more similar to the architecture of plant area than itself later in development whereas plant area in the water-limited treatment block exhibited a genetic architecture similar to the WUE_ratio_ late at the end of the experiment.

Examination of the signed, proportional allelic effects within the greater fixed QTL model indicated that QTL on 2@96, 5@109, 7@99 and 9@34 contribute medium-to-large effects on a majority of the traits examined in both treatment blocks (Fig. 8). The B100 allele associated with QTL on 2@96 and 9@34 both contributed to increased plant size, water loss, WUE_fit_ and WUE_ratio_. The QTL on 2@96 exhibited its greatest influence in the well-watered treatment block whereas the contribution of 9@34 was greater on average in the water-limited treatment block. Both QTL exhibited similar temporal patterns, showing an earlier effect on plant size and WUE_ratio_ but a consistent effect across water loss. Contribution of the B100 allele on 7@99 and 5@109 decrease plant size, water use and the WUE_fit_ traits; the effect of which was greater in well-watered conditions. The magnitude of effects contributed by QTL on 7@99 on plant size decreased through time whereas the effects on water loss and WUE_fit_ peaked after 20 days and decreases slightly thereafter. The 5@109 locus behaves similarly with little temporal variation in plant water use and WUE_fit_. A majority of the other QTL contributed minor effects that became more prominent in one of the two treatment blocks or at a particular developmental time points. Inheriting the B100 allele at QTL on 2@113, 3@48, 4@52, 6@65 and 9@127 increased the values while the B100 allele at the remaining loci (2@11, 5@79, 5@95, 7@34 and 7@53) decreased the value of the traits (Fig. S27).

**Figure 7.**
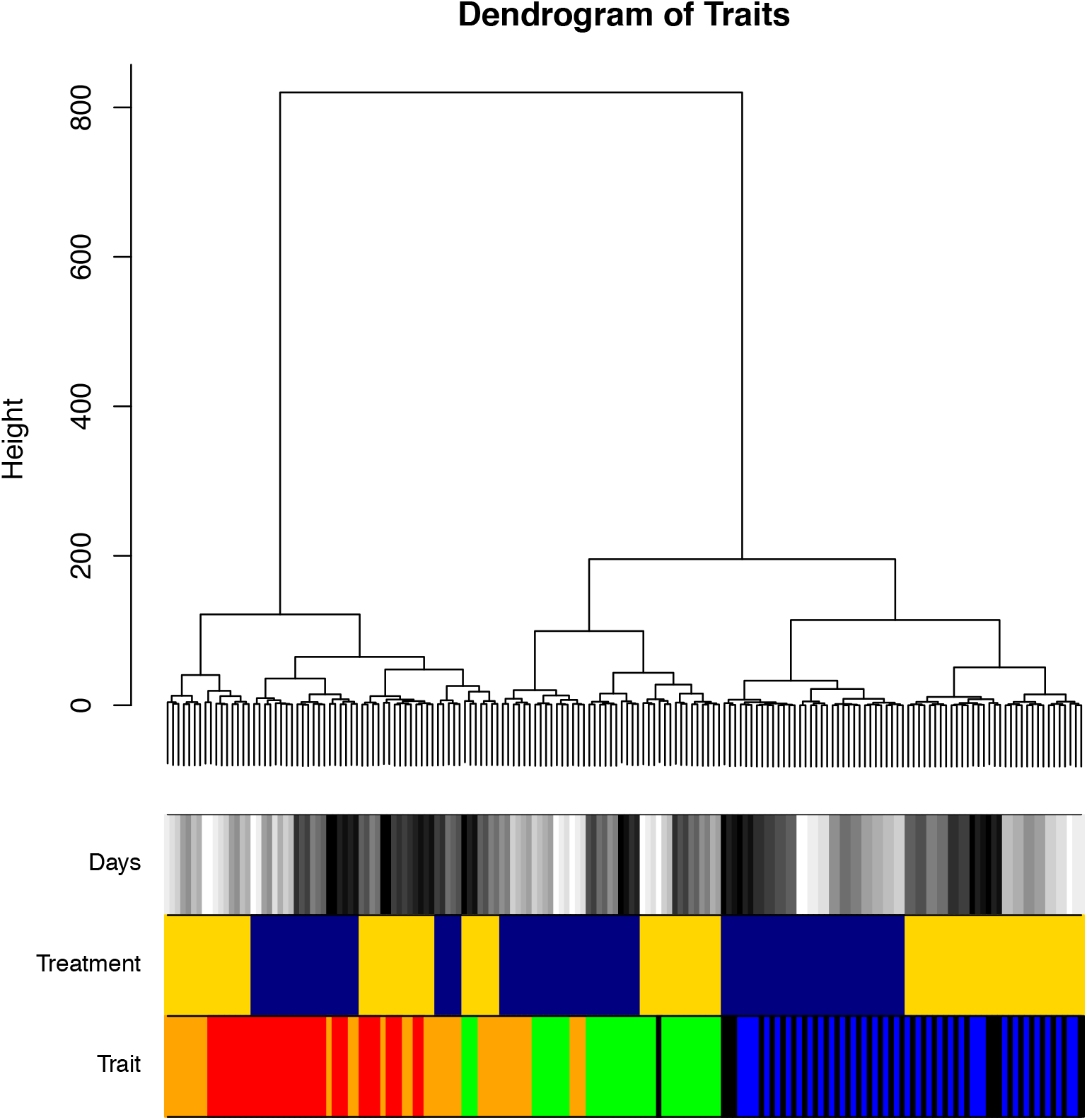
Agglomerative hierarchical clustering defines the relationship between plant size, water use and derived water use efficiency traits. The additive effect size of fourteen common QTLs was calculated across all traits, treatments and developmental time points through hierarchical clustering using Ward’s method. Color bars on the bottom indicate trait (green = plant size, blue = water use, orange = WUE_ratio_, black = WUE_fit_, red = WUE_residual_), treatment block (blue = well-watered, orange = water limited), and days after planting (grey scale values where white represents the trait on day 17 and black indicates the trait on day 33).

**Figure 8.**
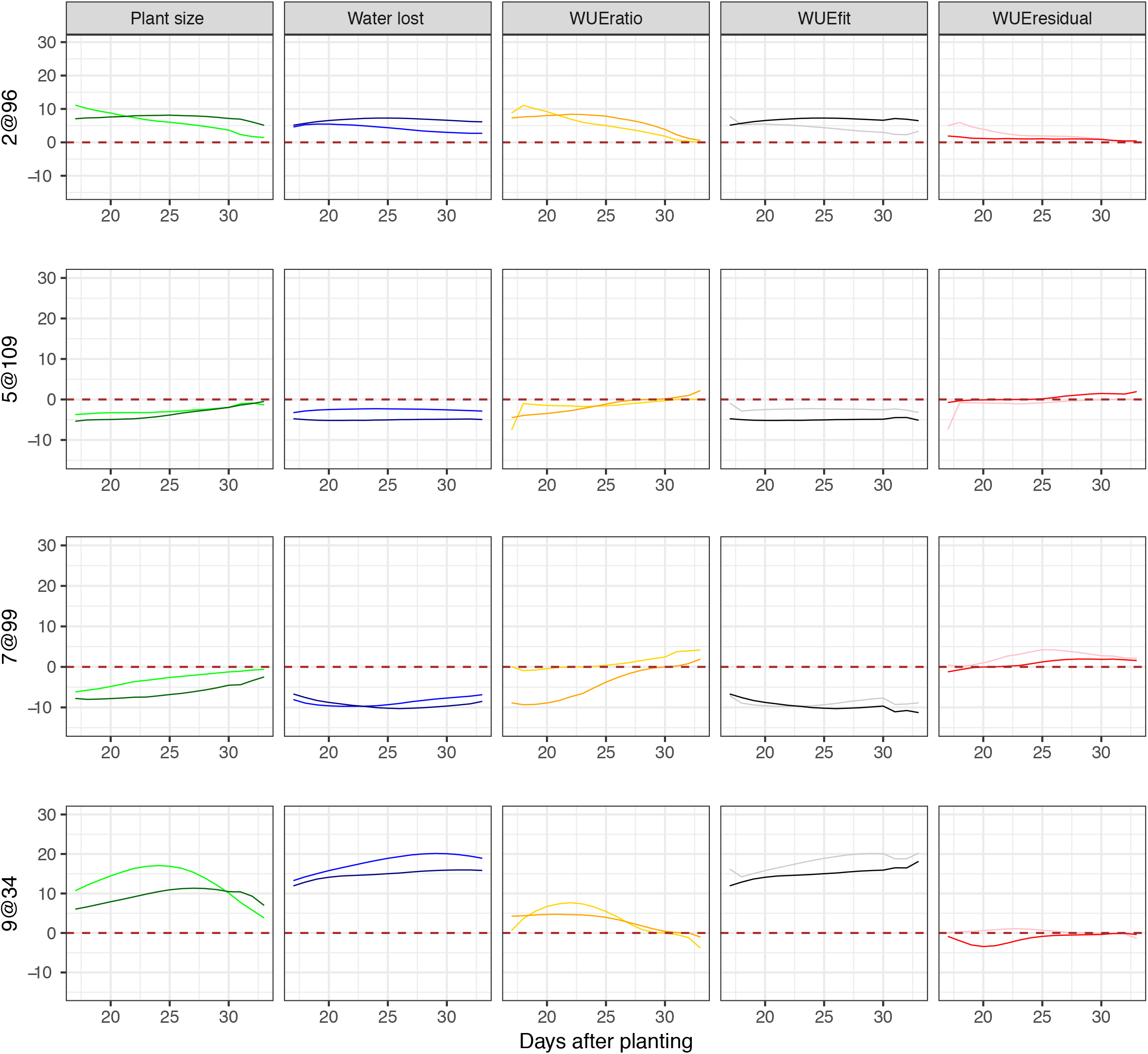
Additive relative effect size of the four major pleiotropic QTL plotted throughout the course of the experiment. A model containing fourteen QTL was fit across traits, treatment blocks and days. The developmental time point (days after planting) is indicated by the x-axis whereas the proportional additive genetic effect size of the B100 allele is plotted along the y-axis. Columns are representative of traits (green = plant size, blue = water use, orange = WUE_ratio_, black = WUE_fit_, red = WUE_residual_) while rows correspond to individual QTL. Shading within the colors denotes treatment block (darker = well-watered, lighter = water-limited).

A majority of the QTL exhibit unidirectional effects across both the well-watered and water-limited treatment blocks although the direction of the effect was largely dependent on the trait (Fig. S28). The exceptions to this trend represent short periods of experimental time at which the relative effect size is near zero within one or both treatment blocks (Fig. 8, Fig. S27).

The proportional contribution of parental alleles towards increased trait values varied between traits, within treatment blocks and throughout plant development. For example, B100 alleles contributed to increased trait values for all traits other than WUE_ratio_ in the water-limited environment and the WUE_residual_ across both treatment blocks (Fig. 9). Alternatively, the contributions of the A10 alleles proportionally increased the WUE_residual_ value early and then again late in plant development relative to those inherited from the B100 parent. The influence of A10 alleles on the WUE_ratio_ was also greater their B100 counterpart under water-limited conditions early in plant development.

## DISCUSSION

**Figure 9.**
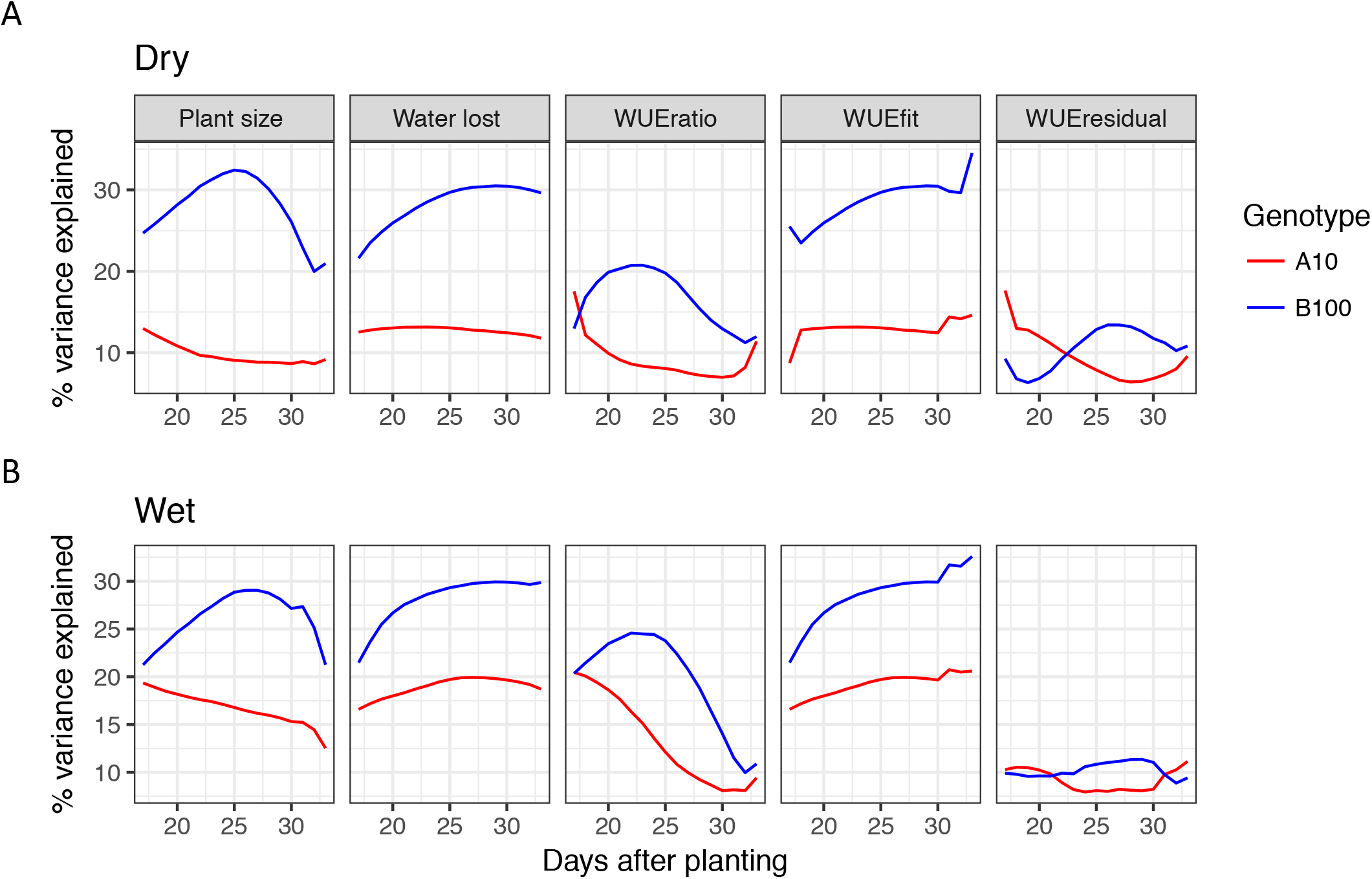
The proportional contribution of parental alleles to increased trait values depend upon trait, environmental water content and plant developmental stage. Alleles derived from the B100 parent contribute a greater proportional of additive genetic variance to plant size, water use and TE model fit in both well-watered and water-limited conditions than their A10 allelic counterparts. Both the WUE ratio and TE model residual traits exhibit dynamic behavior where A10 alleles contribute either greater or close to equal proportions of additive genetic variance early and late in plant development. A) The contribution of parental alleles in the water-limited treatment block. B) The contribution of parental alleles in the water-limited treatment block.

The objectives of this study were to utilize technological advances in high-throughput phenotyping (Chen et al., 2014; Fahlgren et al., 2015; Granier et al., 2006; Pereyra-Irujo et al., 2012; Reuzeau et al., 2006; Sadok et al., 2007; Tisné et al., 2013; Walter et al., 2007) to characterize the genetic architecture of water use efficiency and how this architecture responds to water-limitation in an experimental C_4_ grass model system. Although considerable efforts have been made to characterize these processes in *Arabidopsis thaliana*, C_3_ grass crops and other species (Ruggiero et al., 2017) this represents the first study performed on an annual C_4_ grass RIL population. These efforts enabled us to identify genetic loci that contribute to differential biomass accumulation given water use in a well-watered and water-limited environment. Our findings suggest that the major genetic components associated with plant size, water use and water use efficiency exhibit pleiotropic behavior and that the magnitude of their allelic effects is dependent upon environment and developmental stage. We used two complementary approaches to define traits, and our analysis confirmed that the genetic architecture was similar with both approaches. We show that the loci controlling biomass accumulation can be roughly divided into two groups: those that control the amount of water used to create biomass (WUE_fit_) and those that control how efficiently that water is used (WUE_residual_). The results from this study indicate that alleles from both domesticated foxtail millet and a species representative of its wild progenitor contribute to maximal vegetative biomass yield or water use efficiency grown in environments with different watering regimes. In addition, we highlight aspects of our experimental design and analysis that could be improved in future studies.

### The genetic architecture of plant size, water use, water use efficiency and the relationship between these traits

Within the A10 x B100 *Setaria* RIL population, plant size, water use and the relationship between these two variables are unique polygenic traits whose values are all likely influenced by greater than 10 loci. Four QTL located on 2@96, 5@106, 7@99 and 9@36 exhibit strong pleiotropic influence across this suite of traits, the relative magnitude of each is dependent upon growth environment and developmental time point. Despite strong correlation between plant size and water use we successfully identified genetic architectures distinct to each trait. This was achieved by modeling plant size as a function of water use and examining the resulting values of the model fit (plant size given water use) and deviations from this relationship (residual of plant size given water use). This linear modeling approach has been used much less frequently in the literature (Lopez et al., 2015; Nakhforoosh et al., 2016) than the more commonly used WUE_ratio_ (Adiredjo et al., 2014; Honsdorf et al., 2014; Aparna et al., 2015; Fahlgren et al., 2015; Lopez et al., 2015). While the genetic architectures associated with the WUE_ratio_ and WUE_residual_ in this population are closely related (Fig. 7), WUE_residual_ exhibits substantial heritability and is less correlated with plant size than the WUE_ratio_ (Fig. S6 Fig. S10), making it a more desirable metric.

By examining the model based components of WUE with function valued single marker scan QTL analysis that accounts for multiple hypothesis testing across time points (Kwak et al., 2016), we were able to partition the four major pleiotropic QTL into the genetic components on 2@96, 7@99 and 9@36, which control plant size given water use (WUE_fit_) and those on 2@96 and 5@109 that contribute to deviations from this relationship (WUE_residual_). This result suggests that QTL associated with WUE_fit_ (7@99 and 9@36) potentially control the development of transpiring plant biomass whereas the QTL associated with the WUE_residual_ and WUE_ratio_ (2@96 and 5@109) influence production of non-transpiring tissues or biological processes not directly related to transpiration. This conclusion is in accordance with the results of other studies performed on this population which demonstrate that these loci are largely pleiotropic (Mauro-Herrera and Doust, 2016), although the loci on 2@96 and 5@100 substantially influence plant height (Feldman et al., 2017) and stem biomass, whereas those on 7 and 9 are not associated with the accumulation of stem material (Banan et al., 2017).

Our study also identified many smaller effect QTL which influence biomass, water use and WUE traits. The B100 parental allele contributes substantial positive (3@48, 4@52, 6@65, 9@127) and negative (7@34, 7@53) effects on all traits, whereas QTL on 2@11, 2@113, 5@79 and 5@95 contribute either to plant size/WUE_ratio_/WUE_residual_ ratio to a greater degree than on plant size/water loss/WUE_fit_.

Roughly two thirds of the QTL associated with trait plasticity as a response to water availability (difference or relative difference between treatment blocks) were also identified as being associated with the cumulative traits within both treatment blocks. This observation indicates that in many cases, soil water content influences the temporal dynamics of the allelic effects by differential progression through developmental processes that share similar genetic components (Feldman et al., 2017). This study identifies several QTL (3@48, 7@34 and 9@127) associated with genotype by environment traits which also exhibit significant influence on multiple plant traits within a single treatment block. This provides relatively strong evidence that these QTL have pleiotropic influence on size and water use related traits in an environment specific manner. In contrast, QTL identified only by mapping on the difference or relative difference of the traits between each environment are largely specific to individual traits.

Evidence from this study supports an evolutionary genetic model where the majority of QTL associated with the measured traits exhibit conditional neutrality across both soil water potentials examined. Although all traits other than plant size sometimes exhibit opposite directional effects across treatment blocks, the evidence supporting a model of antagonistic pleiotropy is weak. When identified, QTL exhibiting opposite directional effects within individual treatment blocks were limited to short periods of experimental time and are characterized by negligible relative effects during these time points. The contributions of alleles from both parental lines contribute to increased WUE irrespective of soil water potential, suggesting that neither parent was optimized for WUE. For example, alleles from the A10 parent contribute a greater proportion of additive genetic variance to increased WUE during early development in both well-watered and water-limited environments, (particularly given the WUE_residual_ derivation of WUE) whereas the alleles derived from the B100 parent have greater affect on a majority of the measured traits throughout the time course. The contribution of alleles of both parents to water use efficiency is expected given earlier study performed on the sameplatformwhereparental linesshowedsimilarWUEunderwater-limited conditions (Fahlgren et al., 2015).

### Considerations when measuring plant size, water use and WUE

As observed in many other studies (Chen et al., 2012; Fahlgren et al., 2015; Ge et al., 2016; Golzarian et al., 2011; Honsdorf et al., 2014; Lopez et al., 2015; Parent et al., 2015), relative plant side-view pixel area provided a robust and accurate proximity measurement of plant biomass. Although incorporation of additional plant architectural features can improve estimates of this relationship (Parent et al., 2015), our results indicate that caution should be taken as to not over fit models on ground truth data collected exclusively at the end of the experiment as was performed in this study (Fig. S1).

Automated or manual gravimetric measurement of pot weight has proven to be a reliable estimator of plant transpiration but only if the evaporative loss of moisture from soil can be accounted for. Results presented in this study indicate that inclusion of empty pots (or pots that contain plastic plants (Parent et al., 2015) or fabric wicks (Halperin et al., 2017)) is an appropriate empirical method to estimate the experimental time point at which transpiration contributes meaningfully to total pot evapotranspiration (Coupel-Ledru et al., 2016; Lopez et al., 2015; Pereyra-Irujo et al., 2012). Estimation of evapotranspiration after this critical time point has been effectively used by several other groups to identify and eliminate confounding data points collected early during similar experiments (Vasseur et al., 2014; Coupel-Ledru et al., 2016; Ge et al., 2016). Our findings indicate that subtraction of empty pot weight (as performed by (Pereyra-Irujo et al., 2012; Parent et al., 2015; Coupel-Ledru et al., 2016)) may overcorrect for evaporation at early experimental time points even after the point at which plant transpiration contributes substantially to total pot water loss. Although not applied during this experiment, utilization of plastic covering to shield pots from evaporative moisture loss in combination with the approaches discussed above may improve the ability to unambiguously quantify plant transpiration (Aparna et al., 2015; Coupel-Ledru et al., 2016; Ellsworth et al., 2017; Granier et al., 2006; Halperin et al., 2017; Vasseur et al., 2014). In this study, the contribution of plant biomass to overall pot weight was not accounted for during the estimation of plant water use. Although the contribution of plant biomass to pot weight in most experiments performed using *Arabidopsis thaliana* is negligible (Tisné et al., 2010), plant biomass within this *Setaria* RIL population accounted for 12-18% of total average pot water content by the end of the experiment (Fig. S4). Our inability to account for this growth has the undesirable effect of systematically decreasing the soil water content of larger genotypes, although in practice this small change in soil water potential likely has minimal impact on transpiration dynamics of the plants.

Strong correlation between plant size and water use was observed in spite of the fact that these traits can potentially be controlled by different physiological mechanisms. A similar trend has also been described in experiments designed to study water use efficiency in *Arabidopsis thaliana*, apple and wheat (Lopez et al., 2015; Nakhforoosh et al., 2016; Schoppach et al., 2016; Parent et al., 2015; Vasseur et al., 2014). The magnitude of this correlation is likely inflated in this study due to the large differences in size between parental lines and segregants within the A10 x B100 RIL population. Future studies aimed at investigating the genetic basis of water use efficiency can attenuate this correlation by selecting parental lines of similar size and flowering times that differ in their rates of transpiration within environments of interest.

## CONCLUSIONS

This study leverages recent advances in high-throughput phenotyping and quantitative genetics to identify the genetic loci associated with plant size, water use and water use efficiency in an interspecific RIL population of the model C_4_ grass *Setaria*. Our findings indicate that these traits are highly heritable and largely polygenic, although the effects of four major pleiotropic QTL account for a substantial proportion of the variance observed within each trait. Contribution of parental alleles from both the domesticated and wild progenitor lines contribute to maximization of these characteristics. Overall, the underlying genetic architecture of each of these processes is distinct and substantially influenced by soil water content as well as plant developmental stage. In addition, several aspects of our experimental design which could be improved to obtain a better understanding of the genetic components that underlie plant size, water use and water use efficiency in future high-throughput phenotyping studies.

## REFERENCE

Adiredjo AL, Navaud O, Muños S, Langlade NB, Lamaze T, Grieu P (2014) Genetic Control of Water Use Efficiency and Leaf Carbon Isotope Discrimination in Sunflower (Helianthus annuus L.) Subjected to Two Drought Scenarios. PLoS ONE 9:e101218.

Aparna K, Nepolean T, Srivastsava RK, Kholová J, Rajaram V, Kumar S, Rekha B, Senthilvel S, Hash CT, Vadez V (2015) Quantitative trait loci associated with constitutive traits control water use in pearl millet [*Pennisetum glaucum* (L.) R. Br.]. Plant Biol 17: 1073–1084.

Assouline S, Or D (2013) Plant Water Use Efficiency over Geological Time - Evolution of Leaf Stomata Configurations Affecting Plant Gas Exchange. PLoS ONE8:e67757.

Bacon M (2009) Water Use Efficiency in Plant Biology. John Wiley & Sons, New York, NY.

Banan D, Paul R, Feldman MJ, Holmes M, Schlake H, Baxter I, Leakey ADB (2017) High fidelity detection of crop biomass QTL from low-cost imaging in the field. doi: 10.1101/150144.

Bates D, Mächler M, Bolker B, Walker S (2015) Fitting Linear Mixed-Effects Models Using lme4. J Stat Softw. doi: 10.18637/jss.v067.i01.

Bennetzen JL, Schmutz J, Wang H, Percifield R, Hawkins J, Pontaroli AC, Estep M, Feng L, Vaughn JN, Grimwood J, et al (2012) Reference genome sequence of the model plant Setaria. Nat Biotechnol 30:555–561.

Blatt MR (2000) Cellular Signaling and Volume Control in Stomatal Movements in Plants. Annu Rev Cell Dev Biol 16:221–241.

Blum A (2009) Effective use of water (EUW) and not water-use efficiency (WUE) is the target of crop yield improvement under drought stress. Field Crops Res 112:119–123.

Boutraa T (2010) Improvement of Water Use Efficiency in Irrigated Agriculture: A Review. J Agron 9:1–8.

Boyer JS (1982) Plant Productivity and Environment. Science 218:443–448.

Bozdogan H (1987) Model selection and Akaikes Information Criterion (AIC): The general theory and its analytical extensions. Psychometrika 52:345–370.

Brodribb TJ, Feild TS, Jordan GJ (2007) Leaf Maximum Photosynthetic Rate and Venation Are Linked by Hydraulics. PLANT Physiol 144:1890–1898.

Brodribb TJ, McAdam SAM, Jordan GJ, Feild TS (2009) Evolution of stomatal responsiveness to CO2 and optimization of water-use efficiency among land plants. New Phytol 183:839–847.

Broman KW, Wu H, Sen S, Churchill GA (2003) R/qtl: QTL mapping in experimental crosses. Bioinformatics 19:889–890.

Brutnell TP, Wang L, Swartwood K, Goldschmidt A, Jackson D, Zhu X-G, Kellogg E, Van Eck J (2010) Setaria viridis: A Model for C_4_ Photosynthesis. Plant Cell 22:2537–2544.

Carmo-Silva AE, Francisco A, Powers SJ, Keys AJ, Ascensao L, Parry MAJ, Arrabaca MC (2009) Grasses of different C_4_ subtypes reveal leaf traits related to drought tolerance in their natural habitats: Changes in structure, water potential, and amino acid content. Am J Bot 96:1222–1235.

Chambers JM, Hastie T, eds (1992) Statistical models in S. Wadsworth & Brooks/Cole Advanced Books & Software, Pacific Grove, Calif.

Chaves MM (1991) Effects of Water Deficits on Carbon Assimilation. J Exp Bot 42:1–16.

Chen D, Neumann K, Friedel S, Kilian B, Chen M, Altmann T, Klukas C (2014) Dissecting the Phenotypic Components of Crop Plant Growth and Drought Responses Based on High-Throughput Image Analysis. Plant Cell Online 26:4636–4655.

Chen J, Chang SX, Anyia AO (2012) Quantitative trait loci for water-use efficiency in barley (Hordeum vulgare L.) measured by carbon isotope discrimination under rain-fed conditions on the Canadian Prairies. Theor Appl Genet 125:71–90.

Condon AG (2004) Breeding for high water-use efficiency. J Exp Bot 55:2447–2460.

Condon AG, Richards RA, Rebetzke GJ, Farquhar GD (2002) Improving Intrinsic Water-Use Efficiency and Crop Yield. Crop Sci 42:122–131.

Coupel-Ledru A, Lebon E, Christophe A, Gallo A, Gago P, Pantin F, Doligez A, Simonneau T (2016) Reduced nighttime transpiration is a relevant breeding target for high water-use efficiency in grapevine. Proc Natl Acad Sci 113:8963–8968.

Davies WJ, Bennett MJ (2015) Achieving more crop per drop. Nat Plants 1:15118.

Des Marais DL, Hernandez KM, Juenger TE (2013) Genotype-by-Environment Interaction and Plasticity: Exploring Genomic Responses of Plants to the Abiotic Environment. Annu Rev Ecol Evol Syst 44: 5–29.

Des Marais DL, Razzaque S, Hernandez KM, Garvin DF, Juenger TE (2016) Quantitative trait loci associated with natural diversity in water-use efficiency and response to soil drying in Brachypodium distachyon. Plant Sci 251:2–11.

Devos KM, Wang ZM, Beales J, Sasaki T, Gale MD (1998) Comparative genetic maps of foxtail millet ( Setaria italica) and rice ( Oryza sativa). Theor Appl Genet 96:63–68.

Easlon HM, Nemali KS, Richards JH, Hanson DT, Juenger TE, McKay JK (2014) The physiological basis for genetic variation in water use efficiency and carbon isotope composition in Arabidopsis thaliana. Photosynth Res 119:119–129.

Edwards CE, Ewers BE, McClung CR, Lou P, Weinig C (2012) Quantitative Variation in Water-Use Efficiency across Water Regimes and Its Relationship with Circadian, Vegetative, Reproductive, and Leaf Gas-Exchange Traits. Mol Plant 5:653–668.

Ellsworth PZ, Ellsworth PV, Cousins AB (2017) Relationship of leaf oxygen and carbon isotopic composition with transpiration efficiency in the C_4_ grasses Setaria viridis and Setaria italica. J Exp Bot 68:3513–3528.

Escalona JM, TomàS M, Martorell S, Medrano H, Ribas-Carbo M, Flexas J (2012) Carbon balance in grapevines under different soil water supply: importance of whole plant respiration: Carbon balance in grapevine. Aust J Grape Wine Res 18:308–318.

Evans RG, Sadler EJ (2008) Methods and technologies to improve efficiency of water use: INCREASING WATER USE EFFICIENCIES. Water Resour Res. doi: 10.1029/2007WR006200.

Fahlgren N, Feldman M, Gehan MA, Wilson MS, Shyu C, Bryant DW, Hill ST, McEntee CJ, Warnasooriya SN, Kumar I, et al (2015) A Versatile Phenotyping System and Analytics Platform Reveals Diverse Temporal Responses to Water Availability in Setaria. Mol Plant 8:1520–1535.

Farquhar GD, Hubick KT, Condon AG, Richards RA (1989) Carbon Isotope Fractionation and Plant Water-Use Efficiency. *In* PW Rundel, JR Ehleringer, KA Nagy, eds, Stable Isot. Ecol. Res. Springer New York, New York, NY, pp 21–40.

Feldman MJ, Paul RE, Banan D, Barrett JF, Sebastian J, Yee M-C, Jiang H, Lipka AE, Brutnell TP, Dinneny JR, et al (2017) Time dependent genetic analysis links field and controlled environment phenotypes in the model C_4_ grass Setaria. PLOS Genet 13:e1006841.

Fleury D, Jefferies S, Kuchel H, Langridge P (2010) Genetic and genomic tools to improve drought tolerance in wheat. J Exp Bot 61: 3211–3222.

Flood PJ, Harbinson J, Aarts MGM (2011) Natural genetic variation in plant photosynthesis. Trends Plant Sci 16:327–335.

Franks PJ, Farquhar GD (2006) The Mechanical Diversity of Stomata and Its Significance in Gas-Exchange Control. PLANT Physiol 143:78–87.

Ge Y, Bai G, Stoerger V, Schnable JC (2016) Temporal dynamics of maize plant growth, water use, and leaf water content using automated high throughput RGB and hyperspectral imaging. Comput Electron Agric 127:625–632.

Golzarian MR, Frick RA, Rajendran K, Berger B, Roy S, Tester M, Lun DS (2011) Accurate inference of shoot biomass from high-throughput images of cereal plants. Plant Methods 7:2.

Granier C, Aguirrezabal L, Chenu K, Cookson SJ, Dauzat M, Hamard P, Thioux J-J, Rolland G, Bouchier-Combaud S, Lebaudy A, et al (2006) PHENOPSIS, an automated platform for reproducible phenotyping of plant responses to soil water deficit in *Arabidopsis thaliana* permitted the identification of an accession with low sensitivity to soil water deficit. New Phytol 169: 623–635.

Gregory PJ, George TS (2011) Feeding nine billion: the challenge to sustainable crop production. J Exp Bot 62:5233–5239.

Halperin O, Gebremedhin A, Wallach R, Moshelion M (2017) High-throughput physiological phenotyping and screening system for the characterization of plant-environment interactions. Plant J 89:839–850.

Hamdy A, Ragab R, Scarascia-Mugnozza E (2003) Coping with water scarcity: water saving and increasing water productivity. Irrig Drain 52:3–20.

Hetherington AM, Woodward FI (2003) Theroleof stomata in sensinganddriving environmental change. Nature 424: 901–908.

Holloway-Phillips M-M, Brodribb TJ (2011) Contrasting hydraulic regulation in closely related forage grasses: implications for plant water use. Funct Plant Biol 38:594.

Honsdorf N, March TJ, Berger B, Tester M, Pillen K (2014) High-Throughput Phenotyping to Detect Drought Tolerance QTL in Wild Barley Introgression Lines. PLoS ONE 9:e97047.

Huang P, Shyu C, Coelho CP, Cao Y, Brutnell TP (2016) Setaria viridis as a Model System to Advance Millet Genetics and Genomics. Front Plant Sci. doi: 10.3389/fpls.2016.01781.

Huxman TE, Smith MD, Fay PA, Knapp AK, Shaw MR, Loik ME, Smith SD, Tissue DT, Zak JC, Weltzin JF, et al (2004) Convergence across biomes to a common rain-use efficiency. Nature 429:651–654.

Kenney AM, McKay JK, Richards JH, Juenger TE (2014) Direct and indirect selection on flowering time, water-use efficiency (WUE, δ13C), and WUE plasticity to drought in Arabidopsis thaliana. Ecol Evol 4:4505–4521.

Kwak I-Y, Moore CR, Spalding EP, Broman KW (2016) Mapping Quantitative Trait Loci Underlying Function-Valued Traits Using Functional Principal Component Analysis and Multi-Trait Mapping. G3amp58 Genes Genomes Genetics 6: 79–86.

Lawson T, Blatt MR (2014) Stomatal Size, Speed, and Responsiveness Impact on Photosynthesis and Water Use Efficiency. PLANT Physiol 164:1556–1570.

Lawson T, von Caemmerer S, Baroli I (2010) Photosynthesis and Stomatal Behaviour. *In* UE Lüttge, W Beyschlag, B Büdel, D Francis, eds, Prog. Bot. 72. Springer Berlin Heidelberg, Berlin, Heidelberg, pp 265–304.

Lawson T, Kramer DM, Raines CA (2012) Improving yield by exploiting mechanisms underlying natural variation of photosynthesis. Curr Opin Biotechnol 23:215–220.

Legendre P (2014) lmodel2: Model II Regression. R package version 1.7-2.

Li P, Brutnell TP (2011) Setaria viridis and Setaria italica, model genetic systems for the Panicoid grasses. J Exp Bot 62:3031–3037.

Lopez G, Pallas B, Martinez S, Lauri P-É, Regnard J-L, Durel C-É, Costes E (2015) Genetic Variation of Morphological Traits and Transpiration in an Apple Core Collection under Well-Watered Conditions: Towards the Identification of Morphotypes with High Water Use Efficiency. PLOS ONE 10:e0145540.

Lowry DB, Logan TL, Santuari L, Hardtke CS, Richards JH, DeRose-Wilson LJ, McKay JK, Sen S, Juenger TE (2013) Expression Quantitative Trait Locus Mapping across Water Availability Environments Reveals Contrasting Associations with Genomic Features in Arabidopsis. Plant Cell 25:3266–3279.

Martre P, Cochard H, Durand J-L (2001) Hydraulic architecture and water flow in growing grass tillers (Festuca arundinacea Schreb.). Plant Cell Environ 24:65–76.

Mauro-Herrera M, Doust AN (2016) Development and Genetic Control of Plant Architecture and Biomass in the Panicoid Grass, Setaria. PLOS ONE 11:e0151346.

Mojica JP, Mullen J, Lovell JT, Monroe JG, Paul JR, Oakley CG, McKay JK (2016) Genetics of water use physiology in locally adapted Arabidopsis thaliana. Plant Sci 251:12–22.

Monteith JL (1993) The exchange of water and carbon by crops in a mediterranean climate. IrrigSci. doi: 10.1007/BF00208401.

Morison JI., Baker N., Mullineaux P., Davies W. (2008) Improving water use in crop production. Philos Trans R Soc B Biol Sci 363:639–658.

Nakhforoosh A, Bodewein T, Fiorani F, Bodner G (2016) Identification of Water Use Strategies at Early Growth Stages in Durum Wheat from Shoot Phenotyping and Physiological Measurements. Front Plant Sci. doi: 10.3389/fpls.2016.01155.

Parent B, Shahinnia F, Maphosa L, Berger B, Rabie H, Chalmers K, Kovalchuk A, Langridge P, Fleury D (2015) Combining field performance with controlled environment plant imaging to identify the genetic control of growth and transpiration underlying yield response to water-deficit stress in wheat. J Exp Bot 66:5481–5492.

Pater D, Mullen JL, McKay JK, Schroeder JI (2017) Screening for Natural Variation in Water Use Efficiency Traits in a Diversity Set of Brassica napus L. Identifies Candidate Variants in Photosynthetic Assimilation. Plant Cell Physiol 58:1700–1709.

Penman H, Schofield R (1951) Some physical aspects of assimilation and transpiration. Carbon Dioxide Fixat. Photosynth. 5.

Pereyra-Irujo GA, Gasco ED, Peirone LS, Aguirrezábal LAN (2012) GlyPh: a low-cost platform for phenotyping plant growth and water use. Funct Plant Biol 39:905.

Premachandra GS, Hahn DT, Axtell JD, Joly RJ (1994) Epicuticular wax load and water-use efficiency in bloomless and sparse-bloom mutants of Sorghum bicolor L. Environ Exp Bot 34:293–301.

Reuzeau C, Frankard V, Hatzfeld Y, Sanz A, Van Camp W, Lejeune P, De Wilde C, Lievens K, de Wolf J, Vranken E, etal (2006) Traitmill TM: a functional genomics platform for the phenotypic analysis of cereals. Plant Genet Resour Charact Util 4: 20–24.

Ruggiero A, Punzo P, Landi S, Costa A, Van Oosten M, Grillo S (2017) Improving Plant Water Use Efficiency through Molecular Genetics. Horticulturae 3:31.

Ryan AC, Dodd IC, Rothwell SA, Jones R, Tardieu F, Draye X, Davies WJ (2016) Gravimetric phenotyping of whole plant transpiration responses to atmospheric vapour pressure deficit identifies genotypic variation in water use efficiency. Plant Sci 251:101–109.

Sack L, Holbrook NM (2006) LEAF HYDRAULICS. Annu Rev Plant Biol 57:361–381.

Sadok W, Naudin P, Boussuge B, Muller B, Welcker C, Tardieu F (2007) Leaf growth rate per unit thermal time follows QTL-dependent daily patterns in hundreds of maize lines under naturally fluctuating conditions. Plant Cell Environ 30:135–146.

Saha P, Sade N, Arzani A, Rubio Wilhelmi M del M, Coe KM, Li B, Blumwald E (2016) Effects of abiotic stress on physiological plasticity and water use of Setaria viridis (L.). Plant Sci 251:128–138.

Schoppach R, Taylor JD, Majerus E, Claverie E, Baumann U, Suchecki R, Fleury D, Sadok W (2016) High resolution mapping of traits related to whole-plant transpiration under increasing evaporative demand in wheat. J Exp Bot 67:2847–2860.

Seibt U, Rajabi A, Griffiths H, Berry JA (2008) Carbon isotopes and water use efficiency: sense and sensitivity. Oecologia 155:441–454.

Stanhill G (1986) Water Use Efficiency. Adv. Agron. Elsevier, pp 53–85.

Stewart G, Turnbull M, Schmidt S, Erskine P (1995) 13C Natural Abundance in Plant Communities Along a Rainfall Gradient: a Biological Integrator of Water Availability. Aust J Plant Physiol 22:51.

Tardieu F (2013) Plant response to environmental conditions: assessing potential production, water demand, and negative effects of water deficit. Front Physiol. doi: 10.3389/fphys.2013.00017.

Tisné S, Schmalenbach I, Reymond M, Dauzat M, Pervent M, Vile D, Granier C (2010) Keep on growing under drought: genetic and developmental bases of the response of rosette area using a recombinant inbred line population: Leaf development and drought stress. Plant Cell Environ 33:1875–1887.

Tisné S, Serrand Y, Bach L, Gilbault E, Ben Ameur R, Balasse H, Voisin R, Bouchez D, Durand-Tardif M, Guerche P, et al (2013) Phenoscope: an automated large-scale phenotyping platform offering high spatial homogeneity. Plant J 74:534–544.

Tomás M, Medrano H, Escalona JM, Martorell S, Pou A, Ribas-Carbó M, Flexas J (2014) Variability of water use efficiency in grapevines. Environ Exp Bot 103:148–157.

Vasseur F, Bontpart T, Dauzat M, Granier C, Vile D (2014) Multivariate genetic analysis of plant responses to water deficit and high temperature revealed contrasting adaptive strategies. J Exp Bot 65:6457–6469.

Walter A, Scharr H, Gilmer F, Zierer R, Nagel KA, Ernst M, Wiese A, Virnich O, Christ MM, Uhlig B, et al (2007) Dynamics of seedling growth acclimation towards altered light conditions can be quantified via GROWSCREEN: a setup and procedure designed for rapid optical phenotyping of different plant species. New Phytol 174:447–455.

Wang ZM, Devos KM, Liu CJ, Wang RQ, Gale MD (1998) Construction of RFLP-based maps of foxtail millet, Setaria italica (L.) P. Beauv. Theor Appl Genet.

White TA, Snow VO (2012) A modelling analysis to identify plant traits for enhanced water-use efficiency of pasture. Crop Pasture Sci 63:63.

Winter K, Aranda J, Holtum JAM (2005) Carbon isotope composition and water-use efficiency in plants with crassulacean acid metabolism. Funct Plant Biol 32:381.

Xu Y, This D, Pausch RC, Vonhof WM, Coburn JR, Comstock JP, McCouch SR (2009) Leaf-level water use efficiency determined by carbon isotope discrimination in rice seedlings: genetic variation associated with population structure and QTL mapping. Theor Appl Genet 118:1065–1081.

Zegada-Lizarazu W, Iijima M (2005) Deep Root Water Uptake Ability and Water Use Efficiency of Pearl Millet in Comparison to Other Millet Species. Plant Prod Sci 8:454–460.

Zhou Y, Lambrides CJ, Kearns R, Ye C, Fukai S (2012) Water use, water use efficiency and drought resistance among warm-season turfgrasses in shallow soil profiles. Funct Plant Biol 39:116.

Zhu C, Yang J., Shyu C (2017) Setaria Comes of Age: Meeting Report on the Second International Setaria Genetics Conference. Front Plant Sci. doi: 10.3389/fpls.2017.01562.

